# Study the Effect of Green Method Synthesis Cu- nanoparticles with Ionization Radiation on Lactate Dehydrogenase (LDH)

**DOI:** 10.1101/2024.10.31.621364

**Authors:** Baydaa T. Sih

## Abstract

Copper nanoparticles (Cu-NPs) hold considerable importance in the fields of medicine and pharmacy owing to their pronounced affinity for glucose. These nanoparticles are synthesized utilizing environmentally friendly vitamin C extracted from Iraqi lemons, possessing dimensions of less than 20 nanometers and exhibiting both star and spherical morphologies, which are conducive to efficient cellular penetration.

Optical assessments, including ultraviolet absorption, X-ray diffraction, Fourier-transform infrared spectroscopy (FTIR), and scanning electron microscopy (SEM), have validated that the sizes of Cu-NPs are indeed optimal for applications in biomedicine.

Lactate dehydrogenase (LDH) is an enzymatic protein that plays a pivotal role in cellular respiration. A study conducted on cancer patients undergoing radiotherapy, administered at a total dose of 30 Gy divided into 12 sessions of 2.5 Gy per session, revealed a significant elevation in LDH enzyme levels in both in vivo and in vitro samples, with notable increases up to 7.5 Gy, followed by behavioral alterations observed at dosages of 10 and 15 Gy. The initial doses of irradiation were found to enhance the effectiveness of LDH, thereby facilitating the proliferation of cancer cells.

In an effort to mitigate this effect, Cu-NPs were employed, resulting in a reduction of HDL activity by -36% in the control samples. Nevertheless, exposure to X-ray irradiation modified the characteristics of Cu-NPs, leading to an increase in HDL enzyme values, which may be attributed to alterations in nanostructure or surface interactions.

## Introduction

Copper nanoparticles (Cu-NPs) represent a vital category of nanomaterials within the realms of medicine and pharmacy, attributed to their pronounced affinity for glucose. A multitude of medical investigations has substantiated their therapeutic efficacy. In the course of this study, copper nanoparticles were synthesized utilizing an environmentally sustainable methodology, which incorporated vitamin C (ascorbic acid) extracted from green Iraqi lemons, adhering to a simplified procedural framework. The resultant copper nanoparticles measured less than 20 nanometers in size and exhibited morphologies akin to stars and spheres. Such dimensions and configurations are considered optimal and efficacious within the medical field, facilitating cellular infiltration. Comprehensive optical assessments pertaining to the characterization of Cu-NPs were conducted, specifically encompassing ultraviolet absorption, X-ray diffraction, Fourier-transform infrared spectroscopy, and scanning electron microscopy. Collectively, these analyses corroborated that nanoparticles of diverse dimensions, with a maximum size of 25 nanometers, are ideally suited for application in biomedical research owing to their capacity to traverse cellular membranes without necessitating surface stabilization.

### Lactate Dehydrogenase (LDH)

constitutes one among a collective of enzymes present in blood and various bodily tissues that play a pivotal role in the generation of energy within cells. An elevation in the concentration of LDH within the bloodstream may signify tissue damage, certain malignancies, or alternative pathological conditions. This enzyme is also referred to as lactic acid dehydrogenase and LDH [1,2]. LDH is indispensable during the metabolic conversion of sugars into energy for cellular utilization. It is distributed across numerous organs and tissues throughout the organism, including the liver, heart, pancreas, kidneys, skeletal muscles, lymphatic tissue, and blood cells [3]. LDH is integral to the process of cellular respiration, facilitating the conversion of glucose derived from nutritional sources into bioavailable energy for cellular functions [4]. The LDH enzyme is composed of four subunits, with their specific arrangement varying among different tissues, thus giving rise to distinct isoenzymes. These isoenzymes are designated as LDH-1, LDH-2, LDH-3, LDH-4, and LDH-5, with LDH-1 demonstrating the highest efficiency in converting pyruvate into lactate, whereas LDH-5 is more effective in the reverse reaction.

### Medical Relevance of LDH

The quantification of LDH levels in blood assays serves to evaluate tissue damage, particularly in relation to specific diseases or medical conditions. An increase in LDH levels within the bloodstream may be indicative of tissue damage or cellular necrosis, which can manifest in various circumstances, including:

Tissue damage resulting from physical injuries or contusions.

Hemolytic anemia, wherein red blood cells are subjected to premature destruction.

Hepatic disorders.

Certain cancerous conditions.

Myocardial infarctions and other cardiovascular ailments.

Infections.

Nevertheless, it is important to note that LDH levels are not exclusively indicative of any singular condition. Further diagnostic testing and clinical assessment are necessary to ascertain the precise etiology of elevated LDH enzyme levels [5].

In recent years, nanoparticles (NPs) have gained extensive application in the fields of medicine and pharmacy. NPs are increasingly employed in diagnostic procedures and drug delivery mechanisms. Upon entering the bloodstream, a protein corona develops around the NPs. The dimensions and curvature of NPs are significant characteristics that influence the composition of the protein bound within the corona [6]. Researchers exhibit a keen interest in the versatility of nanoparticle chemistry (e.g., wettability, energy, charge, reactivity) alongside physical properties such as size, shape, and concentration in relation to cellular behavior and protein adsorption [7,8,9]. The introduction of nanoparticles, such as AuNPs, into plasma or serum results in the formation of a hard (h-days) and soft (sec-min) protein corona, thereby establishing a conditioned interface to which cells respond [10,11,12]. Recent investigations reveal critical correlations between NP-protein interactions, immunogenicity, and cytotoxicity [13,14]. Studies have substantiated that the surface curvature of nanoparticles significantly influences the extent of protein adsorption. A diverse array of ligands suggests that protein structure, NP size, composition, and chemistry are paramount in the formation of the protein corona [15,16].

### Applications of Nanotechnology in Medicine

Drug Delivery: The design of nanomedicines aims to selectively target only diseased cells, thereby minimizing adverse effects on healthy cells, particularly in oncological treatment.

Early Diagnosis: Nanoparticles possess heightened sensitivity to specific diseases, rendering them effective as detectors or diagnostic instruments.

Precision Medical Devices: The enhancement of imaging in medical imaging apparatuses such as MRI and X-ray is facilitated by nanotechnology.

### Genetic Research and Therapies

Nanotechnology is paving new avenues in genetic research and enabling the delivery of gene therapies in a precise and targeted manner.

Antibacterial and Antifungal Treatments: The application of nanotechnology serves to eradicate bacteria and fungi, functioning as a means of treatment or sterilization, while concurrently investigating the efficacy of enzymes or hormones within the biological system [17-20].

## Experimental Methods

### 1. Copper Nanoparticles Synthesis

For copper nanoparticles synthesis Cu-NPs, vitamin C (Ascorbic Acid) was prepared from Iraqi green lemons as a reducing agent that can reach a size of 50 nm or less and these sizes, especially below 20 nm, are the most commonly used in the synthesis. Green lemons contain similar amounts of vitamin C (ascorbic acid) to other types of lemons. The ascorbic acid content of green lemons can be similar to that of yellow lemons, typically ranging from 4% to 6% of the lemon’s weight.

#### A-Steps Preparation of Ascorbic Acid

For the isolation of ascorbic acid, green lemons sourced from Hit Farms in Anbar, Iraq were utilized.

1. A kilogram of local green lemons, which underwent washing and cutting while retaining the peel and seeds.
2. Subsequently, 500 milliliters of distilled water were incorporated into the sliced lemons and agitated continuously for a duration of 15 minutes using a high-speed electric blender
3. The resulting concoction was then subjected to freezing at a temperature of -8°C.
4. After being taken out from the freezing unit, the frozen blend was fragmented into small pieces and underwent processing in a specialized grinder for a span of 15 minutes to generate a viscous, chilled liquid.
5. Following this, the mixture underwent filtration through a fine mesh sieve.
6. The resultant extract was transferred into laboratory tubes and subjected to centrifugation, subsequently undergoing filtration with filter paper featuring ultra-fine filtration capabilities of less than 0.5µm specifically utilizing a 12.5cm Whatman (Cytiva) filter.

#### b-Preparation steps

Using (0.06) M copper penta-sulfate and 0.11 M ascorbic acid solution to bromide. All the solution putted in backer (250ml) with a small magnetic stir bar, and a thermometer, solution, the sample was kept on a magnetic stirrer for 24h, 850rpm using (M TOPS (S300HS) med in Sou Korea)). The primitive color of the solution was green, by heating and stirring (45°C) the solution color change to red, then the stagnant appeared with brick red, which is the color of copper. Then it was reddish described. camps were isolated by filtration and cleaning with deionized water. Then wash the solution by centrifuge device at 4500 rpm for 15 min to get residual Cu-NPs, Through the well-known washing method, the precipitate is washed with distilled deionized water three times and the stagnant is filtered on filter paper for (48-72) hours to dry without using an oven and laboratory temperature to avoid the effect of heat on the properties of the resulting Cu-NPs [21].

Cu-NPs with 20-35 nm were prepared by using a green method. Eases Cu-NPs were dissolved by normal saline (to avoid blood hemolysis), in proportions (For each 1 ml of normal saline, 1 µg of Cu-NPs) With the use of a dispersing agent that works to surround the nanoparticles to prevent their adhesion or clumping together (pvp) polyvinyl pyrrolidone at a rate three times the weight of the Nano used(1:3),then placed in a device ultrasonic at a power of 150 watt for 5 minutes.

For measuring the optical density was measured at 340±20 nm (According to BIOLABO company instructions) by using Automatic Semi-biochemist try analyzer BTS-350. XRD,FTIR, UV-absorption and SEM for Cu-NPs was measured.

##### Preparation of Blood Samples

Blood samples were procured from thirty-five individuals diagnosed with breast cancer, with an extraction of 10 ml of venous blood from the antecubital vein. In order to inhibit clot formation, the obtained samples were subsequently placed in containers that contained EDTA.

**Part One:** In Vivo Measurement of LDH Enzyme Activity

This segment is dedicated to the assessment of the LDH enzyme’s activity within the patients’ physiological systems. Blood samples were acquired from the patients at the commencement of their radiotherapy sessions, with prior consent obtained from each individual. Control samples were collected prior to the initiation of radiation therapy, thereby establishing a baseline for subsequent comparison.

Measurements were conducted following each radiation session as outlined below:

**Session 1: 2.5 Gy**

**Session 2: 5 Gy**

**Session 3: 7.5 Gy**

**Session 4: 10 Gy**

**Session 6: 15 Gy (6 MeV)**

Blood samples were extracted four hours subsequent to each session.

**Part Two:** In Vitro Measurement of LDH Enzyme Activity

For the in vitro assessments, control samples from Part One were allocated into ten tubes and categorized into three distinct groups:

**Group One:** Four samples subjected to irradiation via X-ray at doses of 2.5, 5, 7.5, 10, and 15 Gy.

**Group Two:** Five samples administered with 40 µl of Cu-NPs, with one tube designated as a control sample.

**Group Three:** Four samples from Group Two were additionally subjected to X-ray irradiation at the identical doses as delineated in Group One.

The activity of the LDH enzyme was evaluated in a specialized laboratory utilizing a Linear Accelerator X-ray (Primus Mid; Serial No. 3779; Siemens, Germany). The blood samples were irradiated with doses of 5 Gy (divided into four segments of 1.75 Gy each) and 15 Gy (divided into four segments of 3.75 Gy each).

For blood samples obtained from patients who underwent radiation treatment, specimens were collected two hours following the second and fourth sessions. The Linear Accelerator apparatus is situated within the Radiotherapy Department at Al-Amal Hospital for Cancer and is specifically engineered for therapeutic applications. Tubes were positioned in plastic racks beneath the X-ray source, and the exposure time for each sample was automatically calibrated in accordance with the stipulated dose.

**Note: All blood samples were diluted (1 ml of blood to 1.5 ml of diluent) for the purpose of enzyme activity measurement, thereby ensuring uniform conditions across all specimens**.

## Results and Dissection

### 1- Cu-Nanoparticles

#### A- XRD for Cu-NPs

XRD (X-ray diffraction) is used to study the crystal lattice structure of materials and determine the orientation of crystals. In the case of copper nanoparticles (Cu-NPs), XRD analysis usually shows several peaks representing specific locations in the crystal.

XRD peaks represent radial space points that represent specific orientations of atoms within the crystal. In the case of copper, the numbers in parentheses (hkl) express an understanding of the angles and crystallographic directions.

As in figure (1) the peak (1,1,1) represents a point in the crystal that has one parallel axis and two other perpendicular intersections.

**figure (1).**
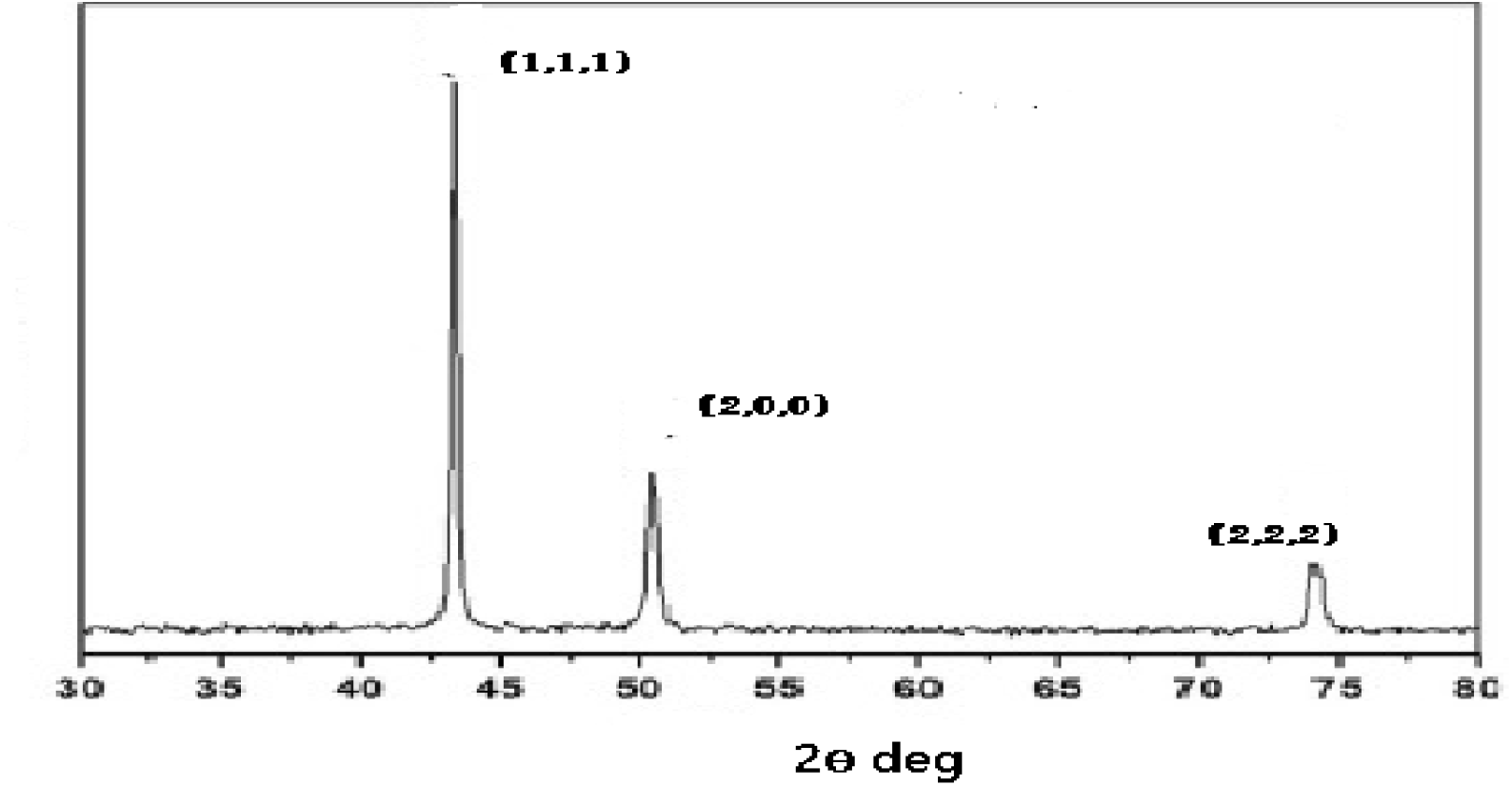
XRD for Cu-NPs.

Peak (2,0,0) means there are two parallel axes.

Peak (2,2,2) expresses the presence of three parallel axes.

If you had an illustration of the XRD pattern of copper nanoparticles, you would notice peaks at known locations of crystallographic orientations such as (1,1,1), (2,0,0), and (2,2,2) for copper.

XRD analysis shows the location and intensity of these peaks and can be used to determine crystal orientations and the size of copper nanoparticles based on the spread and width of the peaks. the lattice distance calculation (d-space) by (using Bragg diffraction) involves determining the length of the Kikuchi Continuous Wave (K.C.W.) which is measured at 1.5406 Å (Å = Angstrom). The break angle (Θ) of the pattern (e.g. 1,1) can be extracted from the graphical representation. Assuming the top 1,1 angle to be 30 degrees, the grid distance is computed accordingly. The process of determining particle size entails the utilization of a specific equation. Essential to this calculation is the knowledge of the line slope (β) derived from the graph’s data. A standard assumption is made for the shape factor (K) at 0.9, serving as an approximate value. The subsequent step involves the actual computation of particle sizes by using Scherrer equation [22].

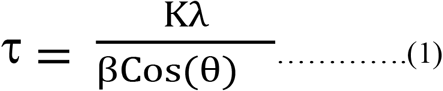

Upon completing the calculations, the following results are obtained: The network distance is determined to be (d = 1.5406) Å, the assumed shape factor (K) is found to be (K = 0.90), and the quantified particle size (utilizing the assumed value for β) is (16.0104) Å.

#### B-UV-Absorption

Absorptivity Measurement: The absorbance of this particular solution is quantified employing a UV spectrometer. The spectrometer illuminates the sample and assesses the extent of light absorbed by the particles within the wavelength range of 200 to 1000 nm.

Data Analysis: Data is systematically gathered and meticulously analyzed to derive an absorption curve that delineates absorption at various wavelengths.

Absorption Curve: The absorption curve illustrates the correlation between wavelength and UV absorption by the sample. The horizontal axis is designated for wavelength (measured in nanometers), while the vertical axis represents absorptivity.

Absorption Peak: The pinnacle of absorption manifests at a distinct wavelength, which is referred to as the absorption peak. In the case of copper nanoparticles, this peak typically occurs around 577 nm, as in figure (2) signifying light absorption at this specific wavelength.

**figure (2).**
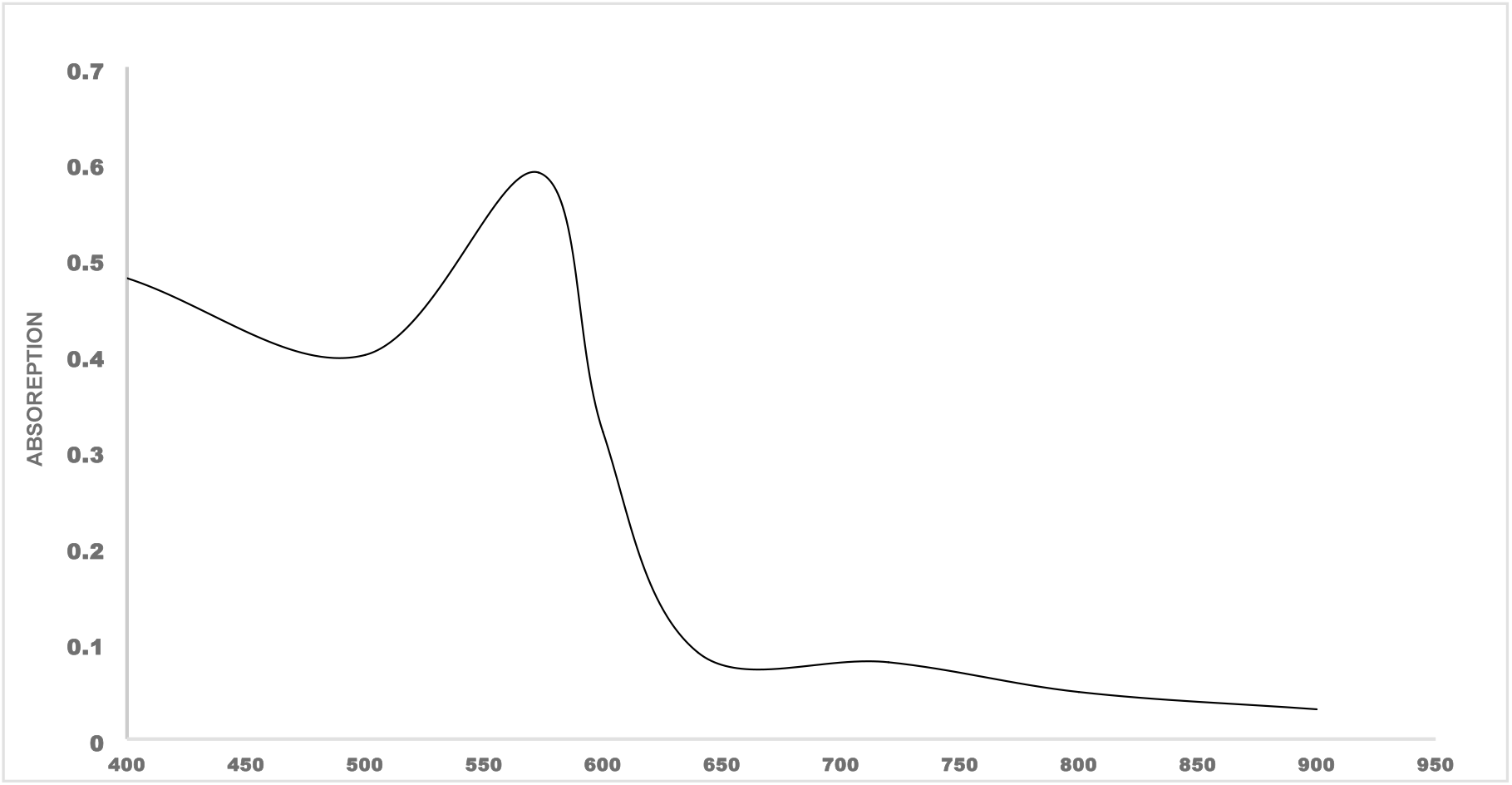
UV-Spectroscopy for Cu-NPs.

Plasmon Surface Resonance: The absorption peak observed at 577 nm signifies the phenomenon of plasmon surface resonance (SPR) associated with copper nanoparticles. This resonance transpires when surface electrons within the particles engage with light, culminating in pronounced absorption at this particular wavelength.

Confirmation of Successful Preparation: The existence of this absorption peak serves as corroborative evidence of the successful synthesis of nanoparticles, as each category of nanoparticles possesses a distinctive absorption spectrum.

#### C- SEM scanning for Cu-NPs

SEM (Scanning Electron Microscopy) analysis is used to see particles at the Nano-level and provide high-resolution images of sample surfaces. When studying copper nanoparticles using SEM, the shape, size and distribution of these particles on the surface can be seen.

Results from an SEM usually contain a set of images that show fine details of the particles. Several pieces of information can be determined from the SEM analysis:

Particle size: Particle sizes can be measured using SEM, which helps in understanding the size distribution and spread of these particles, especially since particle size is very important in working in the medical field, noting that the size below 25 nanometers is the most appropriate for medical work fields, (SEM) used for measuring the size of copper nanoparticles. This device shows different sizes of copper-oxide nanoparticles (14.9, 15.9, 19.7, 20.7, 20.9 and 28.36nm), so the average size is 22.8nm, see figure (3).

**figure (3).**
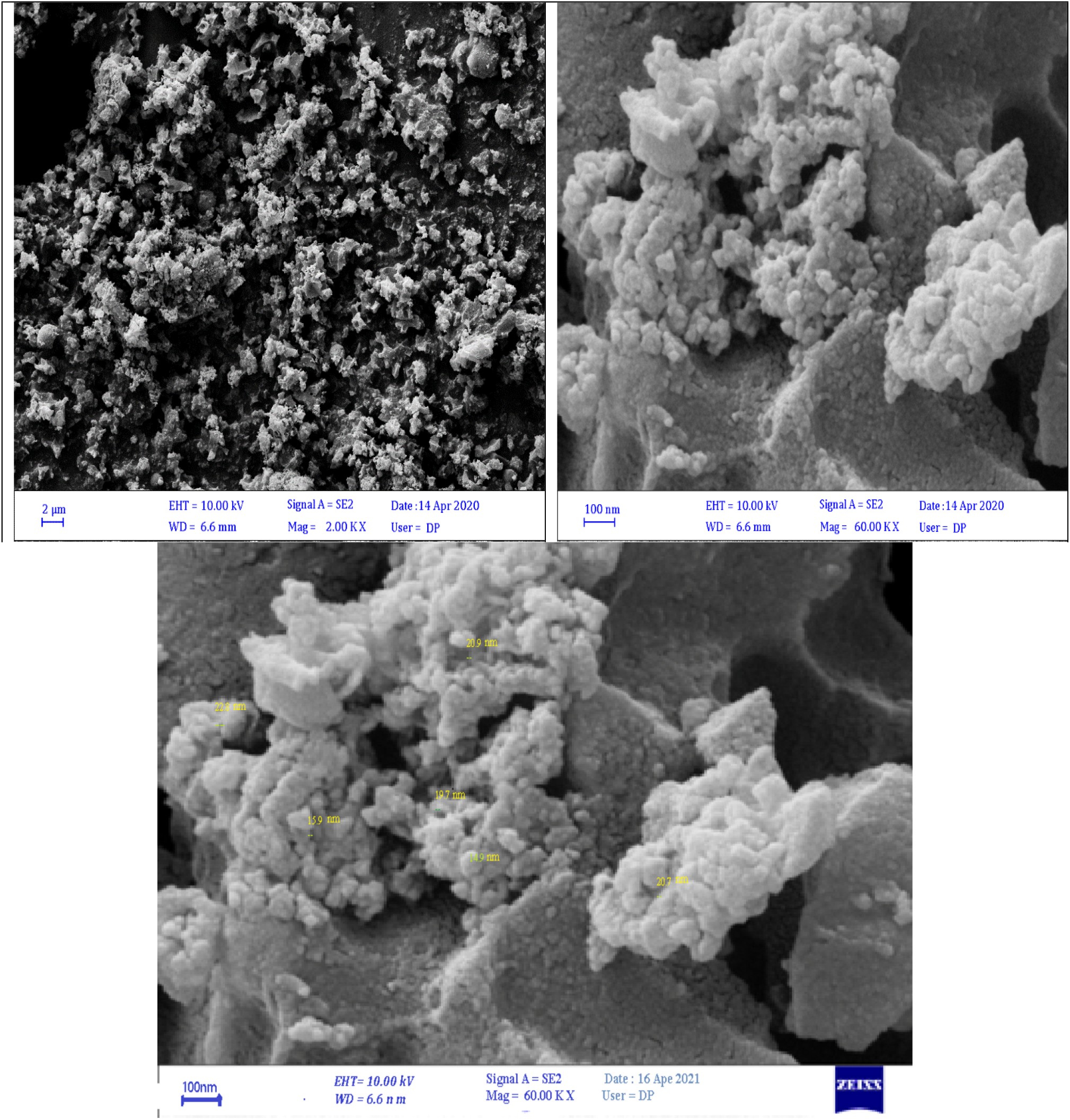
Cu-NPs using SEM technique.

Shape and Structure: SEM shows clear details of the shape of particles, such as whether they are spherical, nearly spherical, or have certain geometric shapes. The surface structure of these particles can also be shown.

Surface distribution: The distribution of particles on the surface and the distances between them can be observed, which gives an idea about the aggregation or surface spread of these particles.

Surface Reactions: SEM may show reactions or changes on the surface of particles, such as possible scattering or impurities.

### 2- Measuring LDH

Experimental results insure that LDH (IU/L) has high value (447±14) U/L) (p-value<0.01) as showed in table (2) in cancer blood samples (including solid tumors), while normal human LDH is between 140 to 280 U/LeAnn important enzyme metabolism of glucose in the cells also increase. Therefore[23,24], it is expected to increase cancer patients blood. This is why this percentage has been shown to be high in the results of patients, and it may be an increase in the cells that secrete this enzyme in the blood. This enzyme indicates increased cell damage and is also an indicator in some cases of the of cancer cells.

**Table (1).**
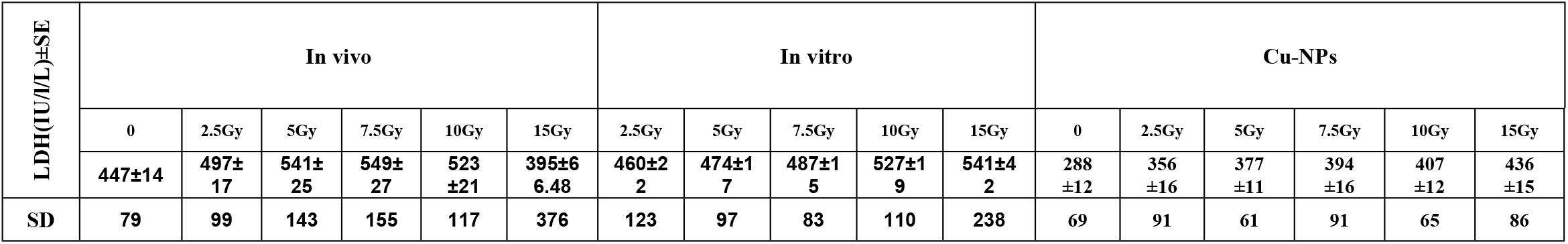
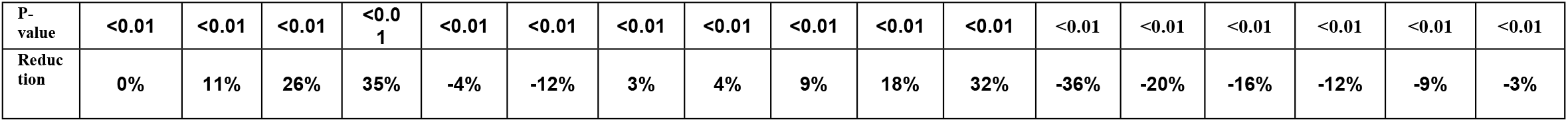
Shows the effect of Cu-NPs and X-ray on LDH activity.

#### The work divided to three parts

a. First part : in this part the change of the enzyme LDH activity with effect of X-ray has been studied for patients after each session of radiotherapy, it’s observed that LDH activity significant increased for the (2.5,5 and 7.5)Gy P-valu ≤ 0.01, but decrease at the dose (10 and 15)Gy. In some clinical studies, researchers confirmed that radiation therapy sessions temporarily increase levels of the enzyme (LDH) in some cases, this increase is often the result of the effect of radiation on the tumor cells and the surrounding tissue, which results in the release of more LDH into the blood. This enzyme is considered to indicate increased cell damage and is also an indicator in some cases of the presence of cancer cells because these cells are voracious consumers of glucose, and since the enzyme is the most important enzyme for glucose metabolism, it is therefore linked to the presence of cells that metabolize more glucose. [25,26]. In 5^th^ session or at 15Gy of radiotherapy almost medical tests results of for patients insure significant decrement (15%) from the control, Radiation therapy targets and kills cancer cells. But the initial doses are not sufficient, so in the beginning it can be concluded that the response of cancer cells is after the dose (7.5Gy). This can lead to a decrease in the number of cancer cells that secrete the LDH enzyme, which results in lower levels of the enzyme in the body. The dose (7.5 and 10)Gy can be considered a dose that improves the patient and reduces the inflammation that accompanies cancer.
b. Part Two: Blood samples taken from Part One patients (for patients before any radiotherapy sessions) were divided into 6 parts, 1.5 ml per tube. Samples were irradiated at the same doses corresponding to radiotherapy doses to study the effect of these doses in vivo. The first three doses (2.5, 5, and 7) Gy were similar in effect but different in values. It can be said that the effect of radiation on cells inside the human body causes a larger area of damage and a change in metabolism compared to outside. In blood samples only blood cells are affected, but in the human body the values of enzymes rise due to the number of affected cells, due to radiation, as it increases Enzyme efficacy in vivo compared to its in vitro increase. After these doses, specifically at (10 and 15) Gray, the behavior differed. In irradiated samples in vitro, enzyme activity increased compared to the control sample, in contrast to what happened in vivo (figure (4)), this is because ionizing radiation increases damage to blood cells with increasing dose in vitro, and this indicates that there is no rebuilding or remodeling, but rather Continuous damage, and this is an indicator of damage to a larger number of blood cells.
c. Third Part: results of treated samples with Cu-NPs showed that significant decreasing in (−36% from the control sample), nanoparticles, especially highly soluble copper, cause the inactivation of the lactate dehydrogenase enzyme. This inactivation may come from a change in some molecules substances or structures present in the blood that causes the cessation of the effectiveness of this enzyme. This opinion is agreed upon by some previous research [27]. It is known that measuring the toxicity of copper nanoparticles causes cell death. It was found that they interact with highly reactive oxygen, and since red blood cells are saturated with
d. oxygen, it is possible to generate reactive oxygen species that cause cell death [28]. In previous research, research has confirmed that the smallest Nano-sizes are the most effective in causing damage in cell membrane, because the cell membranes are saturated with oxygen, and since the hemoglobin in the red blood cell is saturated with oxygen, it is therefore expected that they will be the most affected and exposed to damage. Oxidation: The interactions of copper nanoparticles with the sulfhydryl groups present in enzymes lead to an oxidation and reduction process, which sometimes leads to a loss of the enzyme’s effectiveness. Cu-NPs may be changes in the pH of the enzyme or the combination of a substance HDL with the chemical structure of the enzyme, which disrupts the work of the enzyme or changes its chemical structure, In any case, decreasing LDH enzyme values reduce the growth and proliferation of cancer cells, especially as its known this enzyme is the most important in the process of cellular respiration, by converting lactate into pyruvate [29].
e. Part three: samples treated by Cu-NPs then irradiated by X-ray these samples showed change in the properties of Cu-NPs with X-ray radiation, this change was deduced from the significant increasing 423±19 (P-<0.01) enzyme activity level although it still less than the control by 5%,this increment the surface distribution, crystalline structure, or change in electrical and magnetic properties of the nanoparticles. All of these effects, or more, cause a change in the qualities and properties of the nanoparticles, which finally changes its effect on reducing the effectiveness of the enzyme.

**figure (4).**
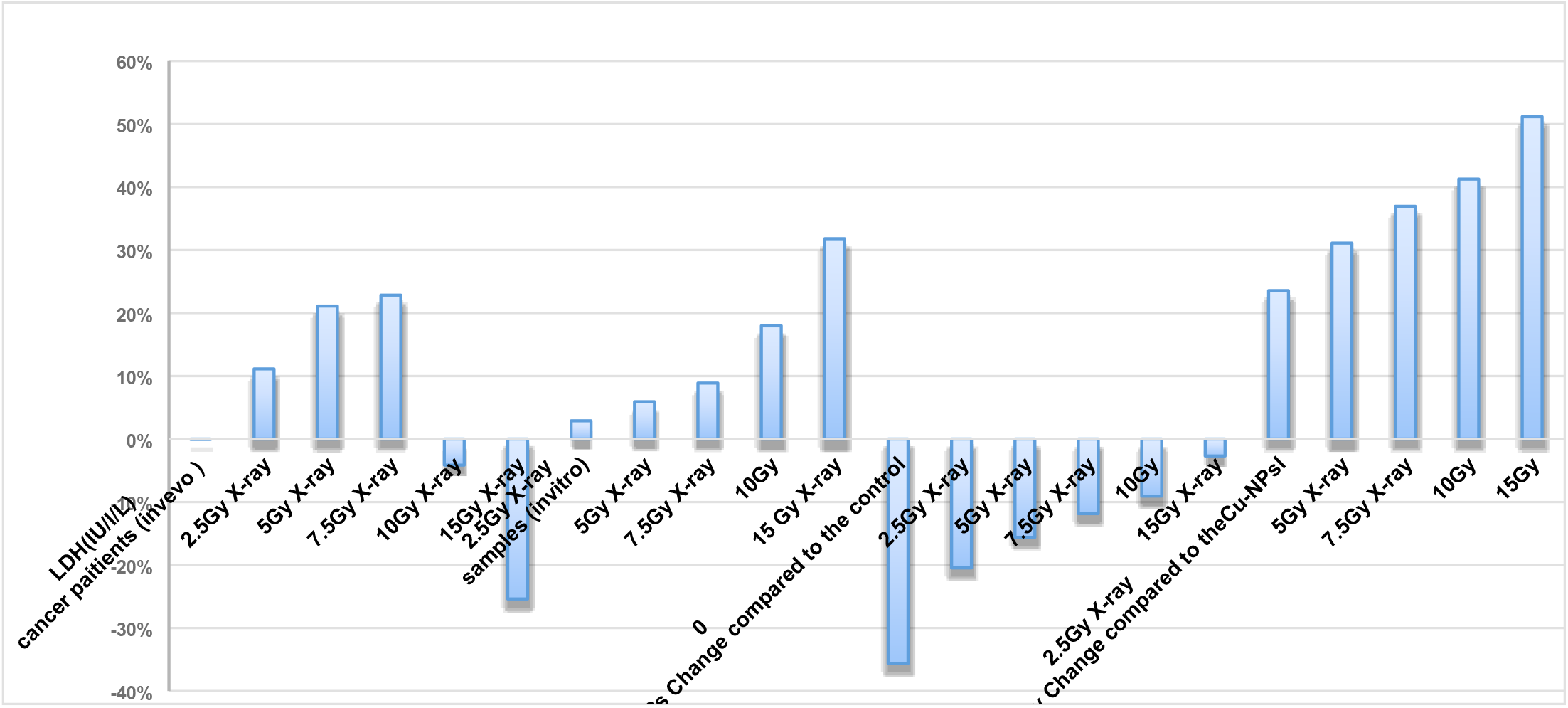
different types of physical effect on LDH enzyme in vivo and in vitro.

In comparison The researcher proved that LDH after radiotherapy may be one of the natural signs of the body’s response to treatment. And in many cases from it is noted that the effectiveness of LDH enzyme increases in irradiated samples entering and(in vivo), but it significantly decreases in samples treated with Cu-NPs. Nevertheless, the effect of the nanoparticles inside the body was not measured due to the inability to administer the nanoparticles to patients. Therefore, the comparison was limited to samples taken from patients receiving radiation treatment and samples irradiated outside the living body, which confirmed the same effect. As for the samples treated with the nanoparticles and then irradiated with the (5 and) dose of X-rays, an increase in the enzyme was observed, but at lower rates than those in the irradiated samples only.

## Conclusions

- LDH enzyme is considered a measure of radiation response in vivo
- Difference in the radiation effects on the effectiveness of LDH enzyme in blood samples in-vitro and in vivo at doses after 7.5Gy of radiotherapy, whereas
- The decrease in enzyme values with the effect of Cu-NPs can be considered an aid in the process of stopping the Krebs cycle in releasing energy to cancer cells and thus stopping their sleep and reproduction.
- The change in the effect of nanoparticles with irradiation on the activity of an enzyme can be considered an indication of a change in the properties of Cu-NPs.

